# Inactivity-mediated molecular adaptations: Insights from a pre-clinical model of physical activity reduction

**DOI:** 10.1101/2024.09.17.613551

**Authors:** Alice Meyer, Nicole Kim, Melissa Nguyen, Monica Misch, Kevin Marmo, Jacob Dowd, Christian Will, Milica Janosevic, Erin J Stephenson

## Abstract

Insufficient physical activity is associated with increased relative risk of cardiometabolic disease and is an independent risk factor for mortality. Experimentally reducing physical activity rapidly induces insulin resistance, impairs glucose handling, and drives metabolic inflexibility. These adaptations manifest during the early stages of physical inactivity, even when energy balance is maintained, suggesting that inactivity-mediated metabolic reprogramming is an early event that precedes changes in body composition. To identify mechanisms that promote metabolic adaptations associated with physical inactivity, we developed a mouse model of physical activity reduction that permits the study of inactivity in animals prior to the onset of overt changes in body composition. Adult mice were randomized into three groups: an inactive control group (standard rodent housing), an active control group (treadmill running 5 d/week for 6-weeks), and an activity reduction group (treadmill running for 4-weeks, followed by 2-weeks of inactivity). Transcriptional profiling of gastrocnemius muscle identified seven transcripts uniquely altered by physical activity reduction compared to the inactive and active control groups. Most identified transcripts had reported functions linked to bioenergetic adaptation. Future studies will provide deeper characterization of the function(s) of each the identified transcripts while also determining how inactivity affects transcriptional regulation in other tissues.

## Introduction

Physical inactivity is one of the leading risk factors for mortality, contributing to >7% of all-cause and cardiovascular deaths annually(1). The global health burden of physical is driven by a relationship between physical inactivity and an increased relative risk of developing a suite of non-communicable diseases such as type-2 diabetes, coronary heart disease, stroke, dementia, and several types of cancer(1, 2). Despite the well-established benefits of regular physical activity on overall health and well-being, approximately one third of the adult population and 81% of adolescents globally do not meet the minimum level of physical activity required for health (i.e., 60 minutes/day of moderate intensity aerobic activities on at least 3 days/week for children and adolescents or 150 minutes/week of moderate intensity aerobic activity for adults)(3–9). In most regions of the world the prevalence of physical inactivity is greatest in girls and women(8–10), while among adults who are classified as being physically inactive, older adults are disproportionately affected(2). Thus, there is a large global population that would benefit from increasing their physical activity levels. Unfortunately, an increasing number of individuals face barriers that prevent them from engaging in exercise for its health benefits(11), thus the prevalence of physical inactivity(1, 12) and inactivity-related disease(13–15) continues to increase.

The past two decades has seen a strong push toward characterizing the physiology of physical inactivity to identify how inactivity drives disease(16–18). This necessary first step has identified a core suite of metabolic adaptations common across multiple models of physical activity reduction. Step reduction(19–24) and bedrest(25–31) protocols in humans and wheel lock(32–39) and cage size reduction(40–43) protocols in animals demonstrate that a reduction in physical activity rapidly induces insulin resistance, impairs glucose handling and drives metabolic inflexibility. These early metabolic adaptations manifest even when energy balance is maintained(25, 27), suggesting inactivity-mediated metabolic reprogramming precedes changes in body composition. However, despite a growing interest in understanding the physiology of physical inactivity, knowledge of the molecular mechanisms that facilitate metabolic adaptations caused by physical inactivity remains limited.

The ability to identify mechanisms responsible for inactivity physiology has been limited, in part, by how physical inactivity is modelled in preclinical settings. Most laboratory animals are housed in environments that restrict activity while concurrently providing free and continuous access to food. Such housing conditions are thought to render animals metabolically morbid making them unsuitable for use as physically active control animals when strategies that further reduce physical activity are employed (i.e., cage size reduction or hind limb casting/unloading)(44). In contrast, while wheel lock models do address the need for animals to be active prior to initiating activity reduction, voluntary wheel running poses its own limitations, such as not being able to easily control the duration and/or intensity of physical activity animals undertake during the active phase of the protocol.

Acknowledging the utility and limitations of existing animal models of physical activity reduction, we developed a treadmill-based rodent model of physical activity reduction. The protocol involves an initial active phase where exercise intensity, duration, and time of day can be tightly controlled, followed by an inactive phase that reduces physical activity without shrinking home cage size or inhibiting hindlimb mobility. Alongside both inactive and active control animals, this manuscript describes this new model of physical activity reduction and reports the transcriptional changes uniquely caused by physical activity reduction in skeletal muscle. We hypothesized that physical activity reduction would cause a unique transcriptional response compared to both active and inactive control animals and that the differentially expressed transcripts associated with physical activity reduction would have functions linked to metabolism.

## Materials and methods

### Animals, housing, diet

Female and male C57BL/6NJ mice (strain #005304) were purchased from The Jackson Laboratory at nine weeks-of age ±3 days. Mice were studied during early adulthood to limit any potentially confounding interactions between physical activity levels and growth and sexual maturation in younger mice, while also limiting any effects of aging on physical activity and other metabolic parameters previously reported to be altered in middle aged or older adult mice(45, 46). Upon arrival at the Midwestern University Animal Facility, mice were housed in a temperature- and humidity-controlled environment (22.2°C, 40% relative humidity) in 7.5”W x 11.5”L x 5”H cages (Ancare #N10) containing corn cob bedding and standardized enrichment (i.e., a plastic tunnel and an iso-BLOX square). Animals were maintained on a 12:12 h light:dark photoperiod cycle and unless otherwise specified, mice had free access to food (Teklad Global #2918; 24% kCal from protein, 18% kCal from fat, 58% kCal from carbohydrate, 3.1 kCal/g) and autoclaved tap water at all times. Throughout the study period, mice were weighed twice weekly using a precision balance under dynamic weighing conditions (Mettler Toledo #MS1602TS). The Midwestern University Institutional Animal Care and Use Committee approved all animal procedures in advance of the study (protocol #3180).

### Treadmill setup and acclimation procedure

All treadmill exercise was conducted on a 5-Lane Touchscreen Convertible Treadmill (Panlab Harvard Apparatus #76-0896) set at a 15° incline. Mice were initially acclimated to the treadmill by placing them on the stationary belt for ∼5 minutes, after which time the belt was started at a speed of 6 m/min. Belt speed was progressively increased in increments of 0.6 m/min every minute until mice were running at a speed of 10 m/min, which they maintained for 5 minutes before being returned to their home cages. This acclimation protocol was repeated up to three times (separate days for each occurrence).

### Physical activity reduction protocol

Ten-week-old mice were randomized into one of three groups (Figure 1a). One group remained inactive (i.e., typical shoe-box style housing with standard enrichment; inactive control group). A second group completed daily treadmill running sessions 5 d/week for 6 weeks (active control group), and a third group completed daily treadmill training 5 d/week for 4 weeks followed by two weeks of inactivity (activity reduction group). Each training session involved a 10-minute warm up period (5 min at 6 m/min, then 5 min at 10 m/min), followed by up to 30 minutes at ∼70% of maximal running speed (predetermined during the prior graded maximal running test, described below; Figure 1b). Duration spent running at 70% of maximal running speed was progressively increased from an initial 15 min to 30 min during the first three weeks of the training program and remained at 30 min/d thereafter (for a total of 40 min running per training session; Figure 1c). To promote training adaptations, running speed was increased by 5% after the first two weeks of training. After 4 weeks, graded maximal running tests were repeated and training speed was adjusted to 70% of the new maximal running speed. All training sessions were conducted between ZT 8-10 h. When necessary, mice were encouraged to run using vocal cues, bursts of compressed air, and/or use of a physical probe to gently tap their hindquarters.

**Figure 1:**
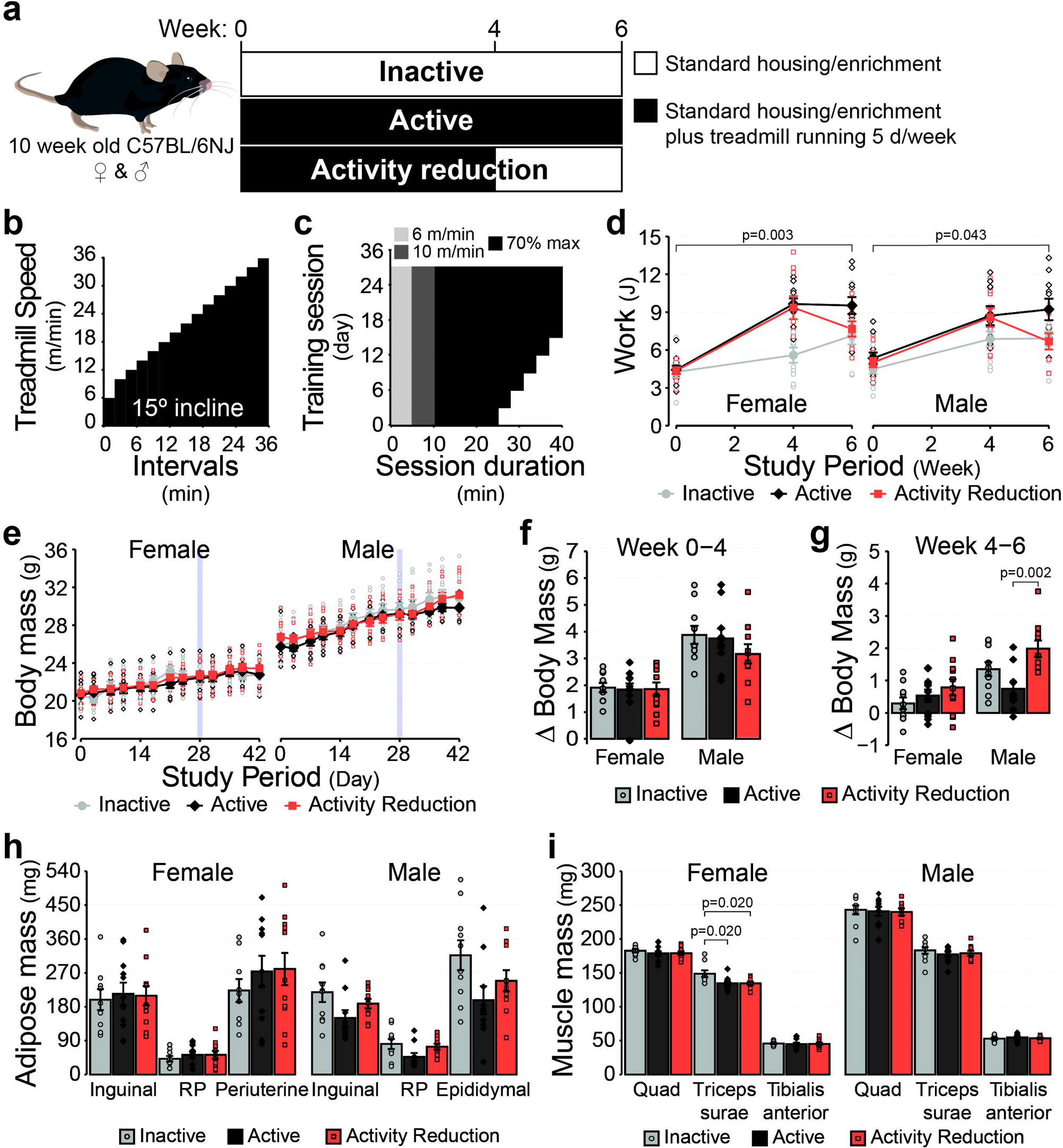
Activity reduction model. (a) Experimental design overview. (b) Overview of the graded maximal running test protocol, which was used to determine training speeds and verify performance adaptations at each stage of the study. (c) Overview of the treadmill training protocol. Training sessions occurred 5 d/week with two consecutive days of rest following every five consecutive days of training. (d) Work (J) calculated for individual mice after each of the graded maximal running tests. (e) Body mass of mice across the period of study. (f) Change in body mass between the start of the study and the end of the fourth week and (g) between the fourth week and the end of the study. (h) Mass of the inguinal, retroperitoneal (RP) and gonadal (periuterine and epididymal) adipose depots at the end of the study. (i) Mass of the quadriceps complex (Quad), triceps surae complex, and tibialis anterior skeletal muscle at the end of the study. Values reported are the mean ± standard error with individual values overlaid for n=30 female (n=10 Inactive, n=11 Active, n=9 Activity Reduction) and n=30 male mice (n=10 Inactive, n=10 Active, n=10 Activity Reduction). Data displayed in panels d and e were compared using linear mixed models with likelihood ratio tests. Data displayed in panels f-h were compared using one-way ANOVA with Tukey tests. Data in panel I was either compared by one-way ANOVA (female quadriceps, male triceps surae, male tibialis anterior), or Kruskal-Wallis tests with Dunn tests and Benjamini and Hochberg corrections (female triceps surae, female tibialis anterior, male quadriceps). Inactive group= light grey circles/bars, Active group = dark grey diamonds/bars, Activity Reduction group = red squares/bars.

### Graded maximal running tests

Maximal running capacity was assessed at the beginning, after 4 weeks, and end of the study using graded maximal running tests. Mice were placed onto a stationary treadmill for up to 5 minutes after which the belt was started at a speed of 6 m/min. Mice maintained this speed for 2 min at which point the speed increased an additional 2 m/min every 2 min until mice reached exhaustion. Exhaustion was defined as the point when a mouse could no longer maintain a running pace that matched the speed of the treadmill and/or they refused to continue running despite gentle encouragement from the person(s) administering the test (as described above). Maximal speed, running distance and running time were recorded. Work during each stage of the test was calculated and summed to determine the total work performed during each test. Work was calculated as: Work J = Force [Body mass kg x 9.18 N] x Vertical Distance [Speed m/sec x Incline (fractional grade) x Time sec].

### Food intake

During the fifth week (i.e., after >1 week of inactivity for the activity reduction group), food intake of individual mice was determined. To facilitate measurement of food intake, mice were moved in pairs into 7.625W” x 15.01”L x 5.13”H cages (Allentown #223581-4) with duel external bottle cage tops (Allentown #228843-1), stainless steel half pocket wire feeders (Allentown #314677) retrofitted with stainless steel dividers, and perforated plastic central dividers that physically separated mice and their food and water access points from one another while retaining many of the social aspects of group housing (Allentown #224520-1). Mice were provided with known amounts of food from which daily consumption was calculated based on the weight of food remaining on subsequent days. Food remaining included the chow pellets retained in the wire feeder basket and, where present, chow pellets and any obvious crumbs recovered from the cage bedding. Energy intake was calculated by multiplying the grams of food eaten by the energy density of the diet (3.1 kCal/g).

### Bomb calorimetry

Stool was collected from cages following the final day of food intake monitoring and stored at -80°C. To prepare samples for calorimetry, stool was dried at 70°C in an oven overnight before being pressed into a single pellet of at least 25 mg using a pellet press. For each sample a 10 cm wire fuse was coiled, placed above the stool pellet and connected to the terminals of the calorimeter (6725 Semimicro Calorimeter, Parr). The bomb chamber was then filled with O_2_ to a pressure of 35 atm and stool pellets were combusted. Energy released during combustion was recorded.

### Tissue collection

Mice were allowed to recover for 48-72 h following the final maximal running test after which they were fasted for 4-6 hours starting at the beginning of the light photoperiod. Mice were anesthetized with isoflurane and blood was collected from the retroorbital sinus using a heparinized capillary and placed immediately on ice. Fasting blood glucose concentrations were measured using a hand-held glucometer (Contour next ONE). Vaginal lavage with sterile PBS was performed on female mice to collect cells for microscopic identification and subsequent determination of estrous stage(47). Following blood collection, anesthetized mice were euthanized by cervical dislocation and tissues rapidly dissected. Right hindlimb muscles were immediately dissected out whole and freeze clamped between stainless steel paddles precooled in liquid nitrogen. Right hindlimb muscles (tibialis anterior and the quadriceps and triceps surae complexes) were carefully dissected out whole and weighed. The left inguinal, gonadal (periuterine or epididymal) and retroperitoneal adipose depots were carefully dissected out and weighed as indices of adiposity.

### Insulin ELISA and HOMA-IR

Fasting blood collected from the retroorbital sinus was allowed to rest on ice for at least 30 min before being centrifuged at 1000 RCF for 20 min at 4°C. Following separation of plasma from the cellular blood components, the plasma was moved into fresh tubes, snap frozen and then stored at -80°C. Frozen plasma was thawed over ice and an ELISA was performed on samples and known standards in triplicate according to the manufacturer’s instructions for a wide-range assay (Crystal Chem #90080). Absorbance was read at 450 nm with background correction for absorbance at 630 nm (Spectramax iD5, Molecular Devices). Fasting plasma insulin concentrations and fasting blood glucose concentrations were used to calculate the Homeostasis Model Assessment of Insulin Resistance (HOMA-IR) using the equation: HOMA-IR = [fasting insulin (µU/mL) x fasting glucose (mmol/L)] / 22.5.

### Glycogen determination in skeletal muscle

The quadriceps muscle complex was powdered while frozen using a liquid nitrogen cooled mortar and pestle. Samples were placed in screw cap vials containing 30% KOH and boiled at 95 °C for 30 minutes with occasional mixing. Samples were then cooled, and 1 M sodium sulfate and 100% ethanol were added. Samples were boiled again for 5 minutes before being centrifuged at 13,000 RCF for 5 minutes at room temperature. After discarding the supernatant, any residual glucose remaining on the glycogen pellet was washed away by resuspending the pellet in water, adding ethanol, and centrifuging the tubes to re-pellet the glycogen. The wash step was repeated three times. Glycogen pellets were allowed to dry at room temperature before being digested overnight at 37°C in 0.3 mg/mL amyloglucosidase prepared in 50 mM sodium acetate, pH 4.8. The next morning, the amount of glucose derived from glycogen was quantified using a commercial glucose kit (Glucose Autokit, Wako diagnostics).

### RNA isolation

RNA was extracted from the entire medial belly of the gastrocnemius muscles. The distribution of fiber types in the mouse gastrocnemius muscle (as determined by myosin-ATPase activity) has previously been described as 5.7-7.6% type-I, 18.1-22.7% type-IIA, 21.7-51.5% type-IID/X, and 18.2-54.4% type IIB fibers(48, 49). Muscles underwent bead homogenization in Trizol. Chloroform was then added to each tube and homogenates were mixed and centrifuged at 4°C to separate the RNA from DNA and protein. RNA was collected from the upper phase and column purified using a commercial kit (PureLink RNA mini kit). RNA concentrations and quality were estimated spectrophotometrically (NanoDrop, ThermoFisher Scientific). RNA quality was determined by denaturing agarose gel electrophoresis(50).

### cDNA library prep and transcriptomics

Aliquots of RNA were sent for commercial sequencing (Azenta). Briefly, RNA integrity was verified by TapeStation (RIN range 8.8-9.8). RNA then underwent Poly(A) selection and cDNA libraries were prepared. Paired end bulk RNA seq was performed (Illumina). Data processing and analysis was completed in house as follows: FASTQ files were imported into Galaxy(51) and sequencing data were trimmed (fastp(52)), aligned to genome assembly GRCm39 (RNA STAR(53)), and reads mapped (featureCounts(54)). Count normalization and differential expression analysis was performed using DESeq2^47^ in R. Transcripts were considered differentially expressed when FDR<0.05 and the fold change was >0.25 in either direction. Gene Set Enrichment Analysis (GSEA) was also performed (fgsea(55)) using the Gene Ontology Biological Processes reference gene set which was downloaded from Molecular Signatures Database(56–58). Enrichment of gene sets is reported where FDR<0.25.

### Statistics

Time course data were analyzed by linear mixed models with likelihood ratio tests and (if applicable) inclusion of fixed factors in the models (see figure legends for details specific to each case), followed by pairwise comparisons using the estimated marginal means to identify differences between individual groups. Single observation data were assessed for homoskedasticity using Levene tests and for normality using Shapiro-Wilk tests. Data that met parametric assumptions were analyzed by either one-way ANOVA with post-hoc Tukey tests or ANCOVA. Non-parametric data were analyzed using Kruskal-Wallis tests with post-hoc Dunn tests. Data wrangling, statistical analyses, and data visualization were performed using R v4.4.1 in R Studio build 764 using the following packages: tidyverse v2.0.0 (59), car v3.1-2(60), rcompanion v2.4.36(61), rstatix v0.7.2(62), lme4 v1.1-35.5(63), and emmeans v1.10.4(64).

## Results

### Running capacity

To ensure that the treadmill training protocol was sufficient to induce a training effect, graded maximal running tests were performed at 0-, 4- and 6-weeks and the amount of work performed during each test was determined (Figure 1d). Group effects were identified for both female (p=0.003) and male mice (p=0.043), with mice in the Active groups performing more overall work than mice in the Inactive groups (p=0.003 for female and p=0.044 for male mice). In contrast, work performed by the Activity Reduction groups was not different from that of the Active (p=0.491 for female and p=0.305) or Inactive groups (p=0.078 for female and p=0.578 for male mice). These findings indicate that not only was the training protocol sufficient to improve performance in both female and male mice, but also that two weeks of inactivity following a four-week training period was sufficient for reversing the effects of regular physical activity on performance.

### Body mass

One goal in developing this model of activity reduction was to make sure data were collected after a period of reduced activity sufficient to ensure the effects being measured weren’t simply the recovery response to the last exercise bout, but also not so long as to be confounded by significant body mass gains. To this end, a two-week period of inactivity was chosen for the Activity Reduction groups and body mass was closely tracked throughout the study. Over the duration of the study, no differences in body mass were observed between groups for either sex (Figure 1e; p=0.895 for female and p=0.505 for male mice). Active male mice tended to weigh less than either the Inactive (-4.5%) or Activity Reduction (-4.3%) groups at the end of 6-weeks, although this effect did not meet a priori assumptions for statistical significance (p=0.220). The amount of body mass gained after the first four weeks also did not differ between groups for either sex (Figure 1f; p=0.974 for female and p=0.335 for male mice). However, between the fourth and sixth weeks, body mass gained by male mice was different between groups (Figure 1g, p=0.003), with male mice in the Activity Reduction group having gained 167.5% more body mass than the Active group during this period (p=0.002). The Inactive males also gained 82.1% more body mass than the Active males; however, this did not meet the a priori threshold for statistical significance (p=0.165). Similarly, although the Activity Reduction males gained 46.9% more body mass than the Inactive males, this difference did not meet requirements for statistical significance either (p=0.143). No differences were detected between groups for body mass gained between weeks four and six for female mice (p=0.276). Together, these data indicate that although two weeks of activity reduction does not lead to a meaningful body mass divergence overall, activity reduction does rapidly reverse the effects of being physically active on suppressing body mass gains in male mice.

### Adipose and skeletal muscle mass

Increases in adiposity and decreases in lean mass have been widely reported to occur in response to physical activity reduction(27, 32, 34, 37, 41). Given these reports and the observation that male mice in the Activity Reduction group experienced catch up weight gain during the two-week inactive phase of the study, we sought to determine whether the Activity Reduction protocol led to changes in the mass of a selection of adipose tissue depots and hind limb skeletal muscles (Figure 1h). For female mice, no differences were observed between groups for the mass of the inguinal, retroperitoneal, or periuterine adipose depots (p=0.744, p=0.481, and p=0.365, respectively). For male mice, the Active group had less adipose mass in their inguinal, retroperitoneal, and epididymal depots when compared to males in the Inactive (-31.4%, -42.0%, and -37.7%, respectively) and Activity Reduction groups (-20.4%, -36.6%, -20.6%, respectively). However, criteria for statistical significance were not quite met for any of the measured depots (p=0.082, p=0.077, and p=0.074, respectively). Since both body mass and the mass of the individual adipose depots were highly variable within individual groups, this comparison was repeated with inclusion of body mass at the time of tissue collection as a covariate in a secondary analysis of the male mice. However, even with body mass included as a covariate, no statistically significant differences were detected between groups for either the inguinal, retroperitoneal, or epididymal depots (p=0.112, p=0.140 and p=0.094, respectively). In both female and male mice, no differences between groups were observed for the mass of the quadriceps complex (p=0.256 and p=0.935, respectively) or the tibialis anterior muscle (p=0.776 and p=0.678). The triceps surae complex was also similar in mass across groups for male mice (p=0.612); however, in female mice that were Active and those that underwent Activity Reduction, the triceps surae complex weighed less than that of the females in the Inactive group (12.9%, p=0.020 and 12.9%, p=0.020, respectively). Although limited by the adipose depots and hindlimb muscles chosen for comparison, and a lack of statistical power given the amount of variability within groups for the adipose depots, these data suggest that two-weeks of Activity Reduction may alter body composition in a sexually dimorphic and tissue-specific manner.

### Energy intake and absorption

Following the observation that male mice experienced catch up body mass gains during the two-week activity reduction period, we sought to determine whether any differences in body mass might be associated with changes in energy intake (Figure 2a) or energy absorption from the diet (Figure 2b). A group effect for energy intake was observed in female mice (p=0.034), but not male mice (p=0.876). Female mice in the Activity Reduction group consumed 32.6% more energy per day than females in the Inactive group (p=0.028). Female mice in the Active group also had 21.5% greater daily energy intake than those in the Inactive group; however, this did not meet a priori criteria for statistical significance (p=0.186). No differences in energy intake were observed between female mice in the Active and Activity Reduction groups (p=0.618). The energy excreted in the stool was also measured as a surrogate measure of energy absorption from the diet (Figure 2b). No differences were observed between groups in either female (p=0.941) or male mice (p=0.438), suggesting that activity reduction had little effect on energy absorption. These findings suggest that in female but not male mice, being active likely regulates energy intake, an effect that persists into the first two weeks of activity reduction. Considering these observations alongside the earlier finding that activity reduction did not lead to differences in body mass or energy absorption in female mice, it’s tempting to extrapolate these results to suggest that energy expenditure was likely greater in female mice in the Active and Activity Reduction groups compared to the Inactive group. In contrast, given that male mice in the Activity Reduction group were experiencing catch up weight gain during the period where energy intake and absorption were measured, and the observation that male mice across all three groups consumed and absorbed similar amounts of energy, we predict that male mice in the Activity Reduction group would have reduced energy expenditure compared to the Active and Inactive groups. Unfortunately, the experimental design used in this study does not allow for the accurate measurement of daily energy expenditure using indirect calorimetry. Thus, future studies using this model of activity reduction will need to consider alternative approaches to tease out the effects of activity reduction on energy expenditure and determine the if there are any sex-specific responses.

**Figure 2:**
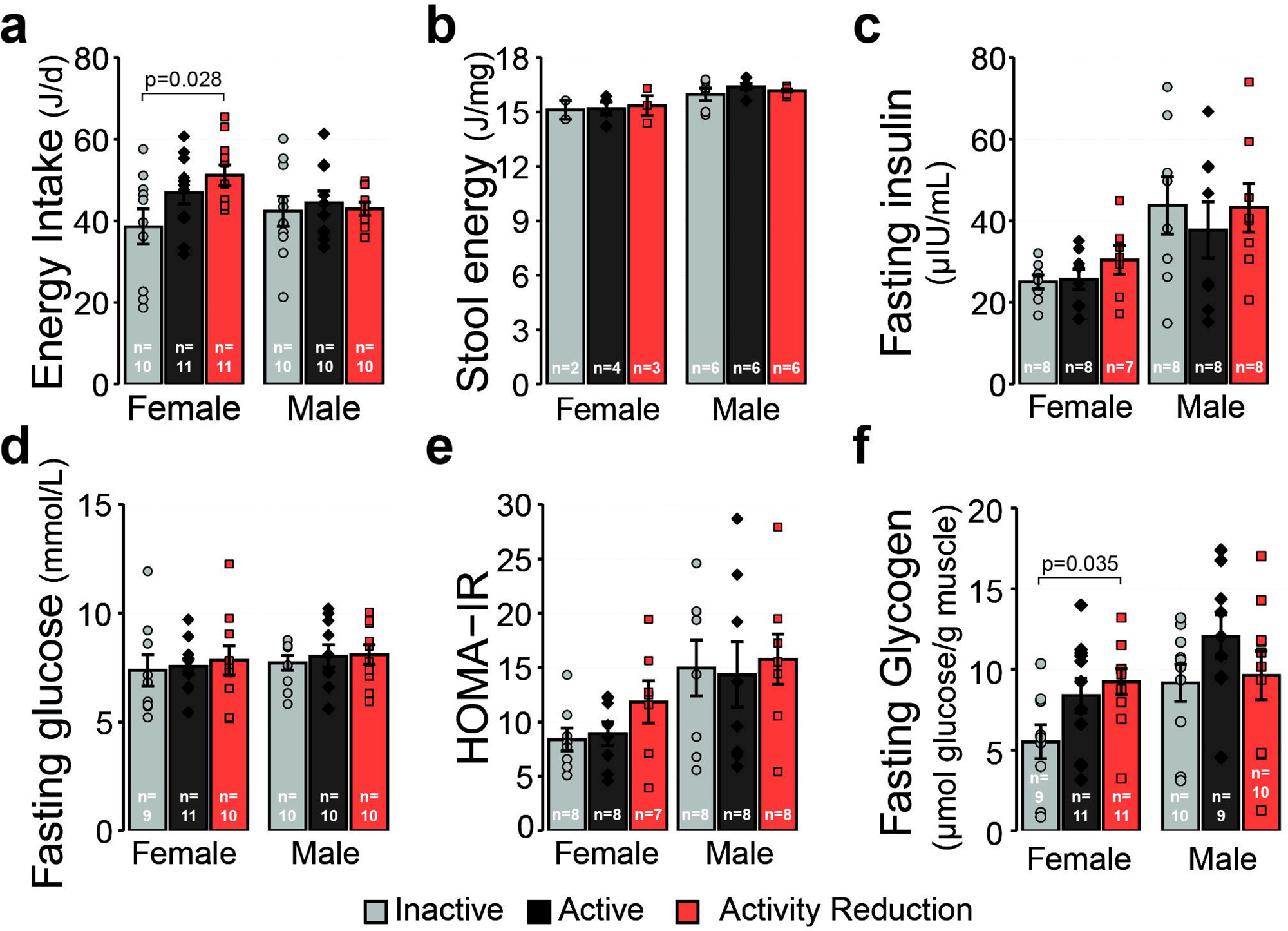
Effect of activity reduction on energy intake, glucose homeostasis and muscle glycogen stores. (a) Average daily energy intake from food in the final week of the study period. (b) Energy content of stool. (c) Fasting plasma insulin and (d) blood glucose concentrations, and (e) HOMA-IR. (f) Glycogen content of the quadriceps muscles. Values reported are the mean ± standard error with individual values overlaid. Data displayed were compared using one-way ANOVA with Tukey tests where appropriate. The number of biological replicates in each experiment is indicated on each individual panel. Inactive group= light grey circles/bars, Active group = dark grey diamonds/bars, Activity Reduction group = red squares/bars.

### Fasting insulin, glucose, and HOMA-IR

To determine whether two-weeks of activity reduction was sufficient to induce hyperinsulinemia, hyperglycemia, insulin resistance, or other disruptions in glucose homeostasis that have been previously reported in other models of activity reduction(20–29, 31, 39, 40, 43, 65), we measured fasting plasma insulin (Figure 3c), fasting blood glucose concentrations (Figure 3d), and calculated the HOMA-IR index (Figure 3e). In both female and male mice, neither fasting insulin concentrations (p=0.304 and p=0.782, respectively) nor fasting glucose concentrations were different between groups (p=0.860 and p=0.808, respectively). HOMA-IR was also not different in male mice (p=0.932); however, there was a tendency for HOMA-IR to be greater in female mice in the Activity Reduction group compared to the Active (32.8%) and Inactive groups (41.2%), although this did not meet the a priori threshold for statistical significance (p=0.197). These findings suggest that two weeks of activity reduction was not sufficient to induce overt disruptions in systemic glucose homeostasis, at least not when animals are in a fasted state.

**Figure 3:**
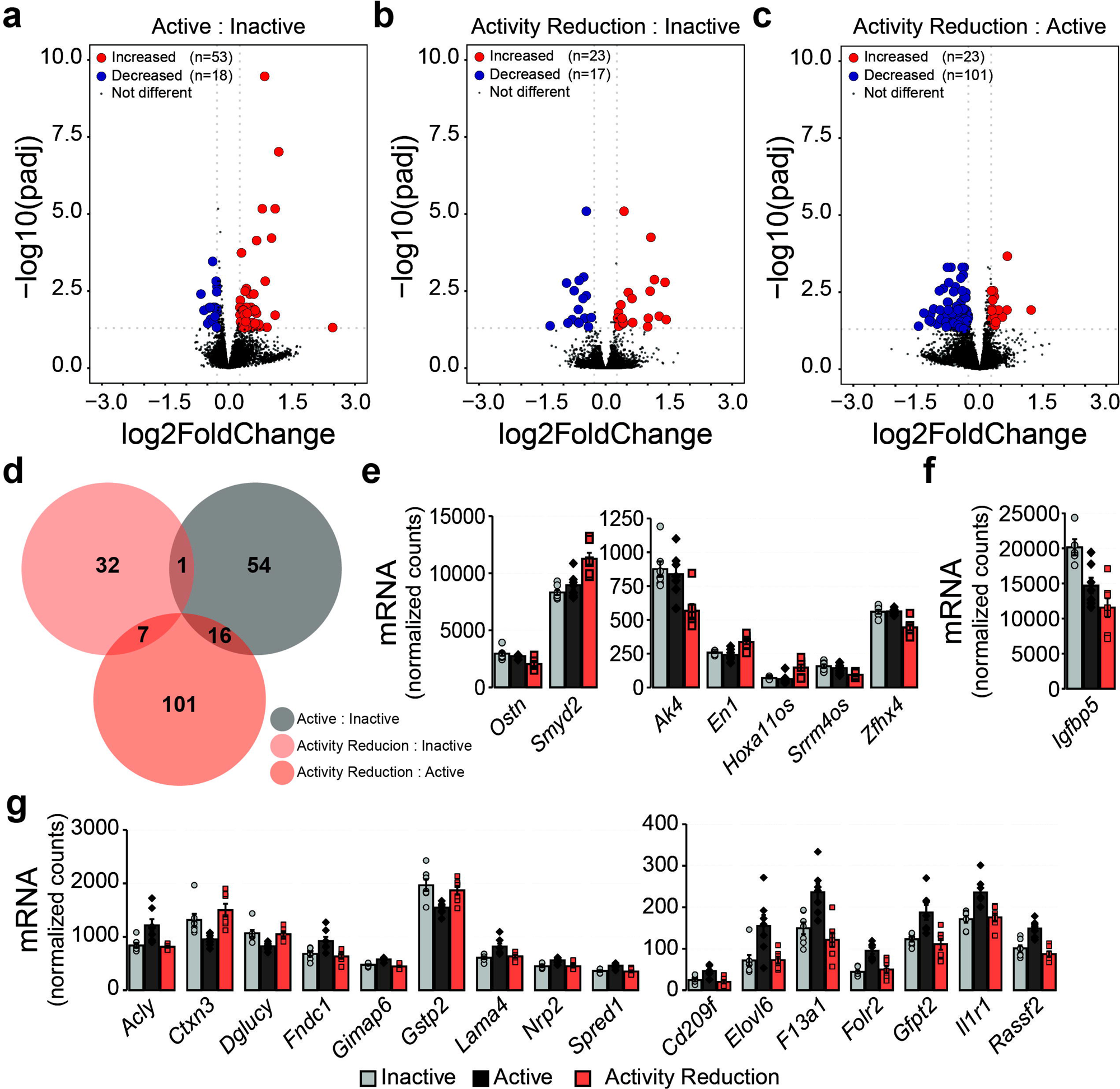
Effect of activity reduction on the skeletal muscle (gastrocnemius) transcriptome. Volcano plots demonstrate the number of up- and down-regulated transcripts in (a) the Active compared to Inactive group, (b) the Activity Reduction compared to Inactive group, and (c) the Activity Reduction compared to Active group. Transcripts were considered differentially expressed when q<0.05 and expression change was greater than or equal to 20% in either direction compared to the Inactive (a and b) or Active (c) groups. Blue = differentially expressed transcripts with reduced expression, red = differentially expressed transcripts with increased expression, grey = transcripts that are either not differentially expressed or are differentially expressed but do not meet the 20% threshold. (d) Venn diagram showing the overlapping and uniquely expressed transcripts in the Active versus Inactive group (grey), the Activity Reduction versus Inactive group (light red), and the Activity Reduction versus Active group (dark red). (e) Expression of transcripts uniquely changed by Activity Reduction compared to both the Inactive and Active groups. (f) *Igfbp5* expression is decreased in both the Active and Activity Reduction groups compared to the Inactive group. (g) Expression of transcripts uniquely changed by Activity compared to the Inactive and Activity Reduction groups. Values reported are the mean ± standard error with individual values overlaid for n=21 female mice (n=7 Inactive, n=7 Active, n=7 Activity Reduction).

### Muscle glycogen concentrations

Since skeletal muscle is a major organ of glucose uptake and storage in the form of glycogen, and a recent study indicated that bed rest in humans leads to increased muscle glycogen storage(65), we sought to determine whether activity reduction in mice also led to changes in the concentration of glycogen in the quadriceps muscle complex (Figure 3f). A group effect for muscle glycogen concentrations was observed for female mice (p=0.037), which was explained by female mice in the Activity Reduction group having 67.6% more muscle glycogen than female mice in the Inactive group (p=0.035). Muscle glycogen concentrations in the Active group were 52.7% greater than those of the Inactive group, but this difference did not meet a priori assumptions for statistical significance (p=0.124). In contrast, although there was a tendency for male mice in the Active group to have greater muscle glycogen concentrations than those in the Inactive (31.5%) or Activity Reduction groups (25.2%), no group effect was detected (p=0.292). These findings indicate that two weeks of physical activity reduction results in sexually dimorphic responses to muscle glycogen concentrations.

### Transcriptional adaptations to activity reduction in skeletal muscle

A major goal of this study was to perform transcriptional profiling in tissues from mice that underwent activity reduction in the hopes of identifying transcripts uniquely changed in response to activity reduction compared to both active and chronically inactive control groups. Bulk RNAseq was performed on RNA isolated from the medial belly of the gastrocnemius muscles from female mice (Figure 3). The Active group had 71 differentially expressed transcripts compared to the Inactive group (Figure 3a), whereas the Activity Reduction group had 40 differentially expressed transcripts compared to the Inactive group (Figure 3b). Compared to the Active group, the Activity Reduction group had 124 differentially expressed transcripts (Figure 3c). After identifying the differentially expressed transcripts in each comparison, differentially expressed transcripts that either overlapped between or were unique to each comparison were determined (Figure 3d). Using this approach, seven transcripts were identified as being uniquely changed in the Activity Reduction group compared to both the Active and Inactive groups (Figure 3e). Of these transcripts, *Smyd2*, *En1*, and *Hoxa11os* all increased in expression following Activity Reduction, whereas *Ostn*, *Ak4*, *Srrm4os*, and *Zfhx4* all decreased in expression. One transcript, *Igfbp5*, was decreased in both the Active and Activity Reduction groups compared to the Inactive group (Figure 3f). Sixteen transcripts were differentially expressed in the Active group compared to both the Inactive and Activity Reduction group (Figure 3g), with the Active group having increased expression of *Acly*, *Cd209f*, *Elovl6*, *F13a1*, *Folr2*, *Fndc1*, *Gfpt2*, *Gimap6*, *Il1r1*, *Lama4*, *Nrp2*, *Rassf2*, and *Spred1* and decreased expression of *Ctxn3*, *Dglucy* and *Gstp2*.

Gene set enrichment analysis was also performed against the Gene Ontology Biological Processes curated gene set (Supplementary Data). Seventy-six gene sets were positively enriched in the Active group compared to the Inactive group. Of these 76 gene sets, all were found to be negatively enriched in the Activity Reduction group when compared to the Active group, suggesting that enrichment of these pathways is dependent on regular physical activity and two weeks of physical activity reduction is sufficient to reverse the activity mediated adaptations. Pathways identified were involved in immune and stress responses, vasculature development and angiogenesis, wound repair, metabolism, and signal transduction. In addition to the pathways found to be positively enriched in response to activity and negatively enriched in response to activity reduction, seven other pathways were identified as being negatively enriched by activity reduction compared to the active group. These include *Endocytosis*, *Import into Cell*, *Vesicle mediated transport*, *Regulation of vesicle mediated transport*, *Regulation of catalytic activity*, *Positive regulation of catalytic activity*, and *Positive regulation of molecular function*. No pathway enrichment was observed when comparing the Activity Reduction group to the Inactive group.

## Discussion

Here, we report an animal model of physical activity reduction that acknowledges the need for animals to be active prior to initiating inactivity, but not so active that they only reflect the activity patterns of those at the higher end of the activity scale. This model offers an alternative to existing models of activity reduction, such as the cage size reduction(40–43) and wheel lock models(33, 66), where investigators are able to control the exercise intensity, exercise duration, and time of day that animals are active. This ensures the activity dose received by both the active control group and the activity reduction group (prior to initiation of inactivity) is consistent and at a volume that can be scaled up or down to reflect different training loads. Since traditional exercise training paradigms designed for increasing performance in humans typically involve exercising at 70% of maximal heart rate or 70% of maximal load, this study employed a training protocol that included a 10 min low speed warm up followed by 30 min running at 70% of maximal running speed 5 d per week. This approach resulted in performance adaptations following the initial four-week training phase that then regressed to levels of the inactive control group following the two-week period of activity reduction. Furthermore, by limiting the activity reduction phase to two weeks and including both active and inactive control groups in the experimental design, the molecular effects of physical activity reduction could be studied without the confounding influence of overt differences in body mass or body composition while ensuring effects specific to activity reduction could be distinguished from the mere reversal of any training-specific adaptations. Limitations of this model also exist. Treadmill exercise is forced exercise which is arguably more stressful to mice compared to voluntary exercise such as wheel running. In the present study we aimed to mitigate the stress response by familiarizing animals to the treadmill ahead of any training sessions or testing of maximal running capacity. Training sessions were also held during the same two-hour window at the end of each light cycle to minimize disruptions during the photoperiod mice are typically least active, and so mice could be familiarized to a routine. Additionally, tissues were collected 48-72 h after all mice had undergone the final maximal running test, thus mice in all three groups received the same treadmill stressor as well as a period of recovery to ensure the acute effects of treadmill exercise and any stress responses associated with it were controlled across all groups. However, whether any of the effects we observed were associated with more chronic adaptations to the stress of forced exercise cannot be ruled out.

Seven transcripts in skeletal muscle were identified as being uniquely changed in response to physical activity reduction. *Smyd2*, *En1*, and *Hoxa11os* were all increased in expression in response to activity reduction. Of these three transcripts, *Smyd2* was by far the most highly expressed. *Smyd2* encodes the lysine methyltransferase SET [Su(Var)3-9, Enhancer-of-zeste and Trithorax] and MYND [Myeloid, Nervy, and DEAF-1] domain containing 2 (SMYD2). SMYD2 monomethylates lysine residues on both histone and non-histone proteins(67) and has largely been studied in the context of tumor biology. Outside of cancer field, SMYD2 has been reported to play a role in regulating titin stability in striated muscle(68, 69). More recently SMYD2 has been found to increase in abundance in the skeletal muscle and livers of hibernating animals during the early stages of torpor(70, 71), providing further evidence that SMYD2 is induced when levels of physical activity are suddenly reduced. Given that SMYD2 has also been linked to the development of metabolic-associated steatotic liver disease(72), we predict that SMYD2 plays a role in regulating metabolic adaptations caused by sudden reductions in bioenergetic demand. Indeed, based on findings in cervical cancer cell lines(73) and our own data (not shown, manuscript forthcoming), *Smyd2* induction in response to reductions in physical activity is likely linked to changes in the way glucose and glucose-products are handled by skeletal muscle.

Although basal expression is much lower in skeletal muscle, *En1*, and *Hoxa11os* were also induced by activity reduction. *En1* encodes engrailed 1 (EN1), which has been shown to bind and regulate the expression of utrophin(74). *En1* has also been linked to pro-fibrotic TGFβ signaling(75, 76), is highly expressed in dopaminergic neurons(77), and has been negatively associated with the thermogenic- and positively associated with the lipogenic capacities of brown adipocytes(78). Less is known about the function of the long non-coding RNA *Hoxa11os*. However, in colon cells, *Hoxa11os* localizes to mitochondria and interacts with several core mitochondrial proteins involved in bioenergetic functions(79). Thus, although the functions of *En1* and *Hoxa11os* in skeletal muscle are currently not well understood, these reports suggest *En1* and/or *Hoxa11os* induction in response to activity reduction could be linked to inactivity-associated mitochondrial adaptations in one or more of the cell populations present in skeletal muscle.

Of the transcripts that were decreased in expression following activity reduction, *Ostn* had the highest basal expression. *Ostn* encodes a the musclin protein (also known as osteocrin), a known myokine that acts as an agonist for natriuretic peptide receptors(80). *Ostn*/musclin induction in skeletal muscle is linked to energy availability, with *Ostn*/musclin expression found to be lower during fasting and higher upon refeeding(80). *Ostn*/musclin is also induced acutely in response to treadmill running(81), indicating that it is exercise-responsive and may contribute to adaptations associated with regular exercise. In line with this notion, when *Ostn*/musclin is absent, impairments in exercise capacity are observed, a finding thought to be due to musclin’s purported influence over mitochondrial biogenesis(81). Conversely, musclin has been identified as a negative regulator of adipocyte thermogenesis(82), and high basal *Ostn*/musclin expression has been reported in animals and people with obesity, insulin resistance, and type-2 diabetes(82–85). In the present study, the expression of *Ostn* was reduced in response to physical activity reduction, as was overall exercise capacity compared to the Active control group. Although merely an association, given previous reports regarding the function of *Ostn*/musclin with regards to exercise tolerance(81), and its role in exercise-mediated cardiac conditioning(86), it is possible that inhibition of *Ostn*/musclin is required for reversal of the effects of regular exercise on functions linked to exercise performance. Whether reductions in *Ostn* expression are associated with changes in circulating musclin concentrations, and whether low *Ostn* expression persists after longer periods of inactivity and/or once significant changes in body composition set in remains to be determined.

*Ak4*, *Srrm4os* and *Zfhx4* were also all decreased in expression in response to activity reduction, although their basal expression was much lower than *Ostn*. *Ak4* encodes adenylate kinase 4 (AK4), a protein that localizes to the mitochondrial matrix(87) and is involved in regulating cellular AMP concentrations and, subsequently, the regulation of AMP-sensitive metabolic enzymes such as the AMP-activated Protein Kinase (i.e., AMPK)(88). Thus, like *Ostn*/musclin, AK4 is also associated with adapting tissues to changes in energy availability. In the present study, physical activity reduction resulted in lower *Ak4* expression in skeletal muscle, an adaptation likely driven by the reduced bioenergetic flux that accompanies physical activity reduction.

Little is known about the function of *Zfhx4*, which encodes zinc finger homeobox 4. *Zfhx4* is highly expressed in developing muscle and proliferating myoblasts, but decreases in expression to very low levels in postnatal skeletal muscles and differentiated myotubes, suggesting its function(s) is/are not critical in tissues with large quantities of postmitotic cells, such as skeletal muscle(89). Indeed, ablation of *Zfhx4* leads to early postnatal lethality linked to respiratory failure, suggesting it plays an important role in early development(90). In colorectal cancer cells, the expression of *Zfhx4* has been computationally associated with the expression of transcripts that promote fatty acid oxidation(91). Thus, considering the known effect of physical inactivity on lowering skeletal muscle fatty acid oxidation(26), and the observation that *Zfhx4* is reduced in expression following physical inactivity, it would not be completely out of line to consider a potential relationship between these processes. Future studies should investigate *Zfhx4* function in postnatal skeletal muscle and determine whether it plays a role in regulating fatty acid oxidation or other functions associated with adapting muscle to changes in bioenergetic demand.

*Srrm4os* encodes serine/arginine repetitive matrix 4, opposite strand. It exhibits low level expression across most tissues(92), although these screens did not include skeletal muscle which is where it was detected in this study and found it to be negatively regulated by physical activity reduction. The function of *Srrm4os* is yet to be determined.

## Conclusion

Seven transcripts in skeletal muscle were identified as being uniquely altered by physical activity reduction. Although the functional contribution of each of the identified transcripts to the physiology of physical inactivity remains to be determined, this study was a necessary first step toward identifying potential mediators of the physiology associated with activity reduction and sets the stage for similar investigations in other tissues. By acknowledging the need for research animals to be more active than typical laboratory housing permits, but not so active the active animals only represent those at the higher end of the activity spectrum, while also ensuring that both active and inactive control groups are included in the experimental design, this work has opened the door to more nuanced investigations into the molecular mechanisms that drive the metabolic impairments associated with physical inactivity. Future studies will provide deeper characterization of the function(s) of each the seven transcripts identified as being uniquely altered by physical activity reduction in skeletal muscle. It is hoped that such next steps will result in identification of molecular targets that can be exploited for the development of therapeutics capable of supporting metabolism for the many people who cannot meet or sustain a level of physical activity required to protect against disease.

## Acknowledgements

We acknowledge past and present members of the Stephenson lab for their contributions, the Midwestern University Core Facility for access to equipment, and members of the Midwestern University community who contributed to discussions that helped shape the final manuscript.

## Author contributions

EJS conceived and designed the research. ES, AM, MM, and KM trained the mice. ES, AM, MM, and MJ collected tissues from the mice. EJS, AM, NK, MN, MM, KM, JD, CW performed experiments. EJS, AM, and NK analyzed data. EJS, AM, and NK interpreted results of experiments. EJS prepared the figures. EJS and NK drafted the manuscript. EJS and AM edited and revised manuscript. All authors approved the submitted version of manuscript.

## Supplementary Material

Supplementary Data includes the raw data used to generate the figures for this manuscript and the output from the differential gene expression and GSEA analyses. See data availability section for instructions on how to access these items.

## Data availability

Sequencing files are available at Gene Expression Omnibus under accession number: GSE281849. Source data and analysis code are available at https://github.com/esteph16/Inactivity-mediated-molecular-adaptations-1.

